# Complete loss of lipoprotein biosynthesis machinery in an obligate intracellular bacterium

**DOI:** 10.64898/2026.10.19.760145

**Authors:** George Winder, Owen Somhorst, Erika Castaneda Sastre, Jack A. Bryant, Jeanne Salje

## Abstract

*Orientia tsutsugamushi* is a cytoplasm-dwelling, obligate intracellular, diderm bacterium with an unusual cell envelope. It lacks lipopolysaccharide and possesses only a minimal peptidoglycan-like cell wall. Here, we report that *O. tsutsugamushi* also lacks the two acyltransferase components of the lipoprotein biosynthesis pathway (Lgt, Lnt) and the entire lipoprotein transport machinery (Lol system). *O. tsutsugamushi* encodes several proteins that are conserved as lipoproteins among other organisms, including Pal and components of the β-barrel assembly machine (BAM) complex, but these lack the canonical lipobox motif, consistent with the absence of lipidation pathways. Instead, we find that *O. tsutsugamushi* Pal compensates for the lack of lipidation by incorporating two predicted amphipathic helices, a feature not observed in Pal homologues from other organisms. These findings suggest that *O. tsutsugamushi* has compensated for a lack of lipoprotein assembly machinery through alternative mechanisms. We speculate on the possible evolutionary advantage of this highly unusual loss of the lipoprotein biosynthesis machinery in the context of an already minimal cell envelope.

**Importance:** Bacterial lipoproteins are nearly universal components of cell envelopes, essential for membrane stability, transport, and pathogenesis. We show that *Orientia tsutsugamushi*, an obligate intracellular bacterium and the cause of scrub typhus, has completely lost the canonical lipoprotein biosynthesis and transport machinery, including lgt, lnt, and the lol system. Despite this, *O. tsutsugamushi* retains several proteins that are normally lipoproteins in other bacteria, including Pal and components of the BAM complex. We demonstrate that Pal instead binds membranes via amphipathic helices, revealing a previously unrecognized mechanism for membrane association. This complete loss of lipoprotein biosynthesis in a diderm bacterium challenges the assumption that lipidation is essential for envelope function and suggests an extreme form of adaptation to a cytoplasmic, intracellular lifestyle. These findings redefine the minimal requirements for a functional bacterial outer membrane and illuminate the evolutionary flexibility of envelope biogenesis in host-adapted microbes.

## Results and Discussion

Bacterial lipoproteins are proteins that are anchored into lipid membranes through covalent attachment of one or more lipid acyl chains^12^. Within the bacterial domain, proteins targeted for lipidation are secreted into the periplasm by Sec or Tat complexes, whereupon lipidation is carried out by three inner membrane proteins. Lgt adds a diacylglycerol moiety to a conserved cysteine residue, LspA cleaves the signal peptide and Lnt adds an additional fatty acid^3^. The lipidated protein is then either retained on the outer surface of the inner membrane or transported to the inner surface of the outer membrane through the Lol transport system comprised of LolA-F. There are two major clades of bacteria. The Gracilicutes contain diderm (Gram-negative) bacteria that typically contain lipoproteins in both their inner and outer membranes. By contrast, the Terrabacteria contain both diderm and monoderm (Gram-positive) organisms and this lineage is characterised by the loss of the Lol machinery and consequent absence of lipoproteins in the outer membrane, where present^4^. Lipoproteins carry out a range of functions in bacteria, and the number can range from approximately 50 to 150 in diderm bacteria and approximately 30 to 80 in monoderm species^5–8^. Important lipoproteins in diderm bacteria such as *Escherichia coli* include Lpp, which covalently bridges the outer membrane to the peptidoglycan layer and provides mechanical stability; Pal, which forms part of the Tol/Pal system to maintain outer membrane stability and coordinate cell division; BamB-E of the BAM complex, which is required for *β*-barrel outer membrane protein processing; and LolB, a component of the lipoprotein transport system which is itself a lipoprotein.

*O. tsutsugamushi* (Ot) is a diderm, obligate intracellular alphaproteobacterium in the order Rickettsiales that resides in *Leptotrombidium* mites, is frequently found in small mammals in endemic regions, and causes the human disease scrub typhus^9^. Ot has a diverse cellular tropism, replicating inside salivary and ovary cells of mites, as well as endothelial, dendritic, monocyte/macrophages and fibroblast cells of mammals. Inside eukaryotic cells, Ot escapes the endolysosomal pathway and replicates directly in the eukaryotic cell cytoplasm^10^. Similarly to other obligate intracellular bacteria, Ot has a reduced genome with associated loss of non-essential genes and a resulting core genome of 657 genes^11^. The cell envelope of Ot is markedly modified compared to that of canonical diderm bacteria, in that it lacks lipopolysaccharide in the outer envelope and builds only a minimal peptidoglycan-like structure for the cell wall^12^. These adaptations are likely at least partially driven by pressure to reduce host cell activation in response to highly immunogenic lipopolysaccharide and peptidoglycan molecules. Similar adaptations have been found in other obligate intracellular bacteria, but these are notably all vacuole-residing organisms^13^. The cytoplasm-dwelling sister genus of Ot, *Rickettsia* spp, retains both lipopolysaccharide and a typical peptidoglycan cell wall^13^. This difference suggests that the cell envelope of Ot has evolved along a distinct trajectory from the closely related *Rickettsia* spp.

Here, we report that Ot completely lacks any detectable homologues of Lgt and Lnt protein acylation machinery, or the Lol export pathway (Fig. 1A), although the signal peptidase LspA is present (Karp_01098). A search for homologues of known lipoproteins revealed the presence of eight proteins: Pal, BamB, BamD, BamE, RlpA, MlaA, AmiD, and MltE. The latter three are not always lipoproteins, including in Alphaproteobacteria whilst RlpA is not a lipoprotein in the sister genus *Rickettsia spp*. (Supp. Fig. 1), so we focussed on the first four. Lipoproteins are acylated at a conserved cysteine residue within a canonical lipobox motif^14^ shown in Fig 1B. This lipobox motif, including the cysteine residue, is conserved in Pal, BamB, BamD and BamE across diverse bacteria including *E. coli*, the model alphaproteobacterium *Caulobacter crescentus*, and a representative from the Ot sister genus *R. prowazekii*. By contrast, no complete lipobox motif containing a cysteine residue could be detected in any of the lipobox homologues of Ot, including Pal, BamB, BamD and BamE. (Fig. 1B and Supp. Fig. 1). However, an incomplete lipobox motif comprising the three lipobox residues immediately upstream of the cysteine residue are present in Pal, BamB and BamE Ot. This may serve as a substrate for the signal peptidase LspA, releasing the proteins into the periplasm following transport across the inner membrane. To further explore the apparent absence of lipoboxes in Ot, the Karp proteome was screened with SignalP 6.0^15^. Sequences predicted as *Sec/SPII* or *Tat/SPll* were collected, and the predicted cleavage sites were inspected for the canonical +1 Cysteine. No candidates contained a +1 Cys, indicating that no canonical lipoproteins were identified. The combined absence of Lgt/Lnt lipidation proteins, Lol transport machinery and complete lipobox motifs indicates that Ot lacks canonical lipoproteins in its cell envelope. A comparison of relevant bacterial species reveals that this complete loss of the lipidation pathway is highly unusual, if not unique to Ot (Fig. 1C, Supp. Table 1).

**Figure 1.**
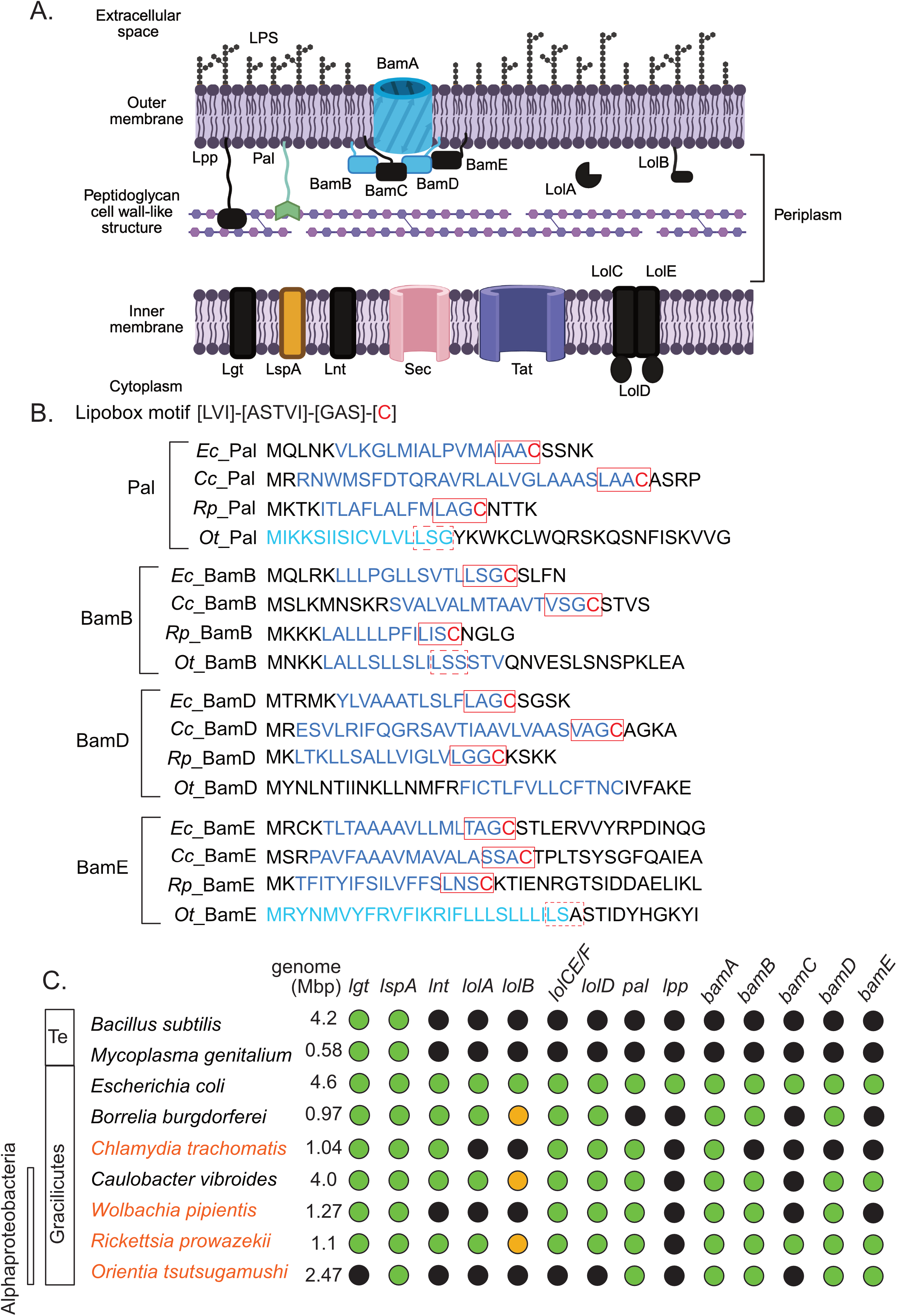
Loss of lipoprotein biosynthesis machinery in Ot. A. Schematic showing the lipoprotein-related components of the cell envelope in Ot, with proteins present in *E. coli* but absent in Ot shown in black. The minimal peptidoglycan-like structure in the Ot periplasm, and lack of lipopolysaccharide (LPS) on the outer membrane, are indicated. Proteins that are normally lipoproteins are shown with lipid anchored extensions. Made using BioRender. B. Signal peptide and lipobox predictions in the four proteins that are normally lipoproteins and present in Ot: Pal, BamB, BamC and BamD. Signal peptides are predicted using SignalP 6.0 (dark blue) and, where that was unsuccessful, DeepTMHMM-1.0 (light blue). Lipobox motifs are predicted using SignalP 6.0 and indicated by a red box (solid line), with the target cysteine residue shown in red. Incomplete lipobox motifs lacking a cysteine residue are shown with a dotted red line. The canonical lipobox motif is shown at the top of the panel. C. Analysis of the presence and absence of genes involved in lipoprotein biosynthesis as well as key lipoproteins, across a range of relevant bacterial species. Green indicates gene presence, black indicates absence, and orange indicates ambiguity. Explanation for these analyses is provided in Supp. Table 1. Te = terrabacteria. The genome size is given and obligate intracellular bacteria are shown in orange.

The lack of lipoprotein biosynthesis capability in Ot leads to three possible outcomes for the proteins: they are not membrane associated, they use an alternative lipoprotein biosynthesis pathway, or they are anchored to the membrane via an alternative mechanism. To explore these possibilities, we examined the lipoprotein Pal. Canonical Pal is comprised of a globular peptidoglycan-binding domain that is anchored to the inner leaflet of the outer membrane through lipidation^16^. A comparison of the predicted three-dimensional structure of Ot_Pal with *Rickettsia prowazekii* Pal revealed similarity of the peptidoglycan-binding domains combined with a unique extension of the Ot_Pal N-terminal domain (Fig. 2A) that we did not observe in any Pal structures from other organisms. Inspection of this extended N-terminal region revealed the presence of two predicted amphipathic helices (Fig. 2B, C). A liposome pelleting assay was used to test whether Ot_Pal binds to lipid membranes and whether the putative amphipathic helices are required. This showed that full length Ot_Pal pellets with liposomes made using *E. coli* total lipid extract, whilst an N-terminal truncation lacking both predicted amphipathic helices does not (Fig. 2C, D). Both purified proteins were shown to be folded using a Thermal Shift Assay.

**Figure 2.**
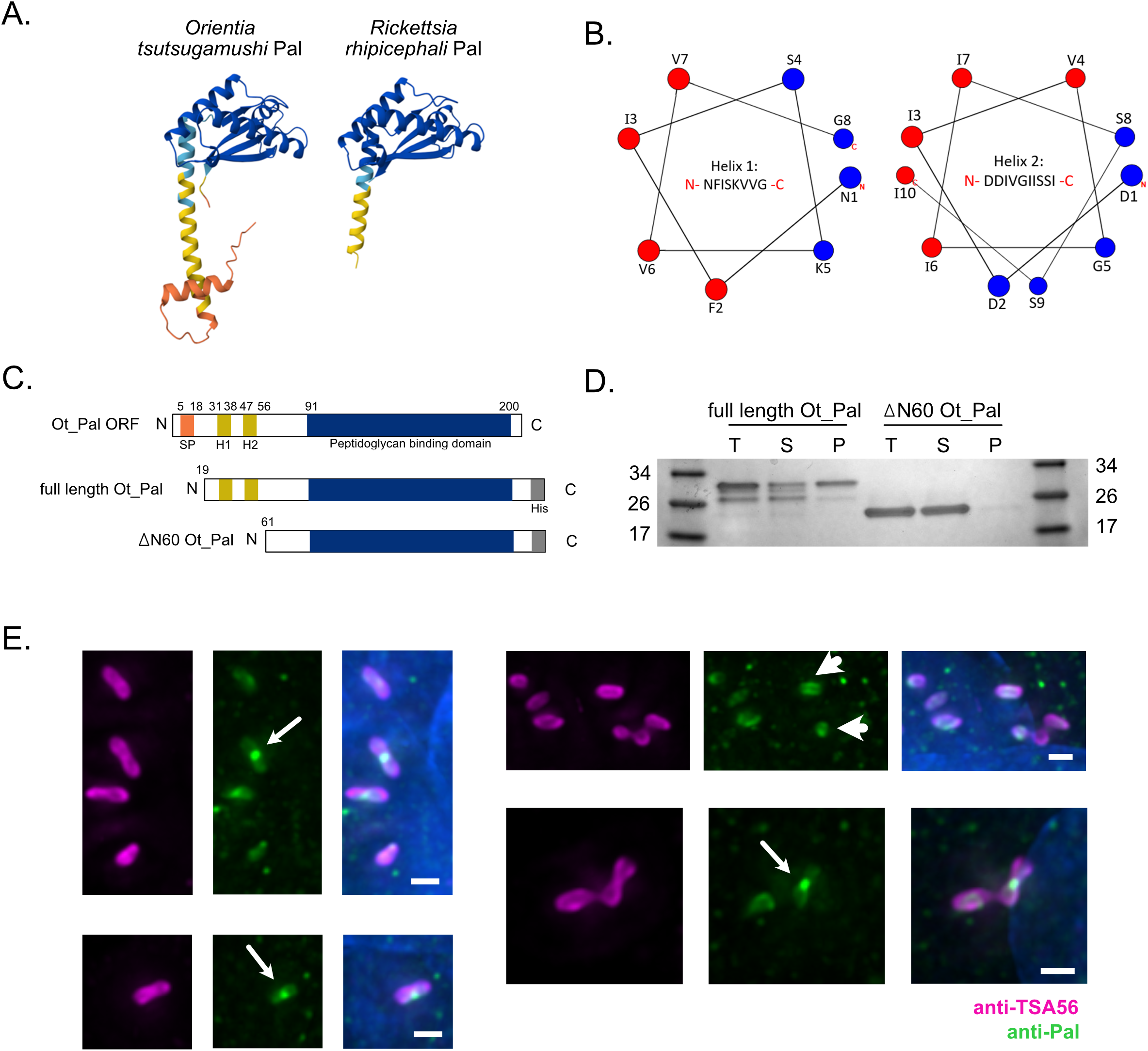
Ot_Pal binds directly to lipid membranes. A. Three-dimensional model of Pal lacking signal peptides from Ot and *Rickettsia prowazekii* predicted using Alphafold 3 showing an N-terminal extension unique to Ot_Pal. B. Hydrophobicity plot of alpha helices at positions 31-38 and 47-56 showing that both form predicted amphipathic helices with distinct hydrophobic (red) and hydrophilic (blue) faces. C. Overview of the Ot_Pal open reading frame, and the full length and ΔN60 constructs used in liposome pelleting assays. D. Results of liposome pelleting assay using full length and ΔN60 constructs described in C. T = total protein, S = supernatant, P = pellet. Full length Pal pellets with liposomes, ΔN60 Pal does not. E. Immunofluorescence microscopy images showing localisation of Pal in Ot cells. Ot_Pal is localised to the cell periphery (arrowheads) and the septum of dividing bacteria (arrows). Bacteria were labelled using an antibody against Ot_Pal (green) and an antibody against the major surface protein TSA56 (magenta). Scale bar = 10 µm.

While we have demonstrated that Ot_Pal associates with liposomes *in* vitro, it remained unknown whether Ot_Pal was associated with the cell envelope in Ot. A polyclonal antibody was generated against Ot_Pal and used to assess the localisation of Pal in Ot cells. Immunofluorescence microscopy revealed that this protein localises to the cell periphery and the septum of dividing cells (Fig. 2E). Whilst the resolution is not sufficient to distinguish whether this is embedded in the inner or outer membrane of the bacterium, it is consistent with membrane association in Ot.

In summary, we report that the cytoplasmic obligate intracellular diderm bacterium Ot completely lacks the canonical lipoprotein biosynthesis machinery and recruits Pal to lipid membranes through predicted amphipathic helices. It remains unknown whether and how the other proteins that lack lipobox motifs are recruited to the membranes. Similar amphipathic helices were not readily predicted in BamB and BamD, which may be recruited to membranes through interactions with the rest of the BAM complex or may not associate with membranes at all. This work raises the question of why Ot has dispensed with protein acylation, a process universally conserved in the bacterial domain. It is unlikely to result simply from genome reduction, as there are many smaller genomes that retain this pathway. It is possible that the loss evolved to reduce immunogenicity through activation of patter recognition receptors, however this step is not taken by any other host-associated bacteria. It is notable that the absence of lipidation is observed within the context of a loss of lipopolysaccharide and reduced peptidoglycan, suggesting an overall weakening of the cell envelope. This may be required for interactions with components of the host cytoplasm that remain to be discovered.

## Materials and Methods

### Mammalian cell culture

Human retinal pigment epithelial (RPE-1, ATCC CRL-4000) and mouse fibroblast clone 929 (L929, ATCC CCL1) cell lines were maintained as monolayers in T25 or T75 cell culture flasks in Dulbecco’s Modified Eagle Medium (Gibco) supplemented with 10% (v/v) heat-inactivated fetal bovine serum (Gibco), at 37 °C in a humidified atmosphere with 5% CO_2_. At 80-90% confluence, cells were either infected with Ot or sub-cultured using trypsin (Gibco).

### Culture of Ot in host cells

Ot strain TA686 was grown in L929 cells under the same conditions as uninfected cells. Cells were infected by adding Ot that had previously been isolated from L929 cells. Culture medium was replaced 3-24 h after infection.

For standard bacterial purification, culture medium was discarded, and flasks were washed with phosphate buffered saline (PBS). Infected cells were harvested with cell scrapers (Corning) in 1 mL of medium, pelleted by centrifugation at 800 x *g* for 5 min at 4 °C, and resuspended in 1 mL of PBS. Bacteria were released using a bead mill homogeniser (Fisherbrand) on setting 5 for 1 min, with 0.1-0.15 mm glass beads (BioSpec). Centrifugation at 800 x *g* for 5 min at 4 °C removed cell debris and beads. If required for subsequent steps, bacteria were pelleted by centrifugation at 14,000 x *g* for 10 min at 4 °C. For long term storage, bacteria were resuspended in SPG buffer and stored at −80 °C.

### Plasmids

Plasmids containing constructs for the liposome pelleting assay were ordered from Twist Bioscience after design using SnapGene 7.0.3 (www.snapgene.com). Plasmids were codon optimised for *E. coli* expression under the IPTG-inducible T7 promoter and contained a kanamycin resistance gene.

Pal sequences were obtained from the Ot strain Karp Pal sequence (GenBank LS398548, locus 261246-261860).

### Protein purification

Plasmids were transformed into *E. coli* strains BL21 and DH5α (New England Biolabs) for protein expression and long-term storage, respectively. *E. coli* strain BL21 carrying plasmids pGW1, pGW2 were cultured overnight from the −80 °C stock then sub-cultured in 1L of LB broth with 50 µg/mL kanamycin and grown to optical density at 600 nm (OD_600_) of 0.4-0.6. Gene expression was induced with 400 µM IPTG (Fluorochem) at 16 °C for 16 h with shaking (180 rpm), then bacteria were pelleted by centrifugation at 3,500 x *g* for 3 min at 4 °C and stored at −80 °C.

Pelleted bacteria were thawed at 37 °C and immediately resuspended at 4 °C in 40 mL of lysis buffer (0.5% v/v Triton X-100, 300mM NaCl, 30mM imidazole, 50mM HEPES). Cells were then pulse sonicated at 4 °C for a total of 10 min at 20 kHz (60% amplitude, pulse: 15 s on, 45 s off), followed by centrifugation at 100,000 x *g* for 40 min at 4 °C, using the Optima XL-100 K ultracentrifuge, SW 32 Ti swinging-bucket rotor and 38.5 mL tubes (Beckman Coulter). Soluble fractions were collected at 4 °C and applied to a column (Pierce) containing Ni-NTA resin (HisPur), 5 mL at a time. Insoluble fractions were resuspended in 40 mL of ultra-pure water. Columns were washed with 25 mL of wash buffer (500mM NaCl, 50mM imidazole, 50mM HEPES) and proteins were eluted with 5 mL of elution buffer (500mM NaCl, 500mM imidazole, 50mM HEPES). For immunoblotting, 80 µL samples were collected from each fraction, heated at 95 °C for 5 min in sodium dodecyl sulfate (SDS) loading buffer and stored at −20 °C. Elution fractions were concentrated 50-fold by repeated centrifugation in protein concentrators (Pierce) at 4000 x *g* for 3-5 min and resuspended in storage buffer (150mM NaCl, 50mM HEPES), with a sample also taken after 5-fold concentration. The concentrated eluate was stored at −80 °C.

Protein concentrations were estimated using a BCA Protein Assay kit (Pierce) according to manufacturer’s instructions.

### Liposome pelleting assay

Liposome pelleting assays were carried out using *E. coli* total lipid extract (Avanti). Lipids were dissolved in ethanol and vortexed, before evaporation of the solvent using a nitrogen stream. The dried lipid film was hydrated using storage buffer and vortexed briefly, allowing multilamellar liposomes to form. Liposomes, at 0.8 mM in 50 µL, were incubated with 1 µM protein for 1 h at room temperature with rocking, then pelleted by centrifugation at 20,000 x *g* for 10 min. The supernatant was collected, and the pellet was washed with 50 µL of storage buffer. The centrifugation was repeated, and the supernatant was discarded. The pellet was resuspended in 50 µL of storage buffer.

Next, 15 µL of SDS loading buffer was added to the resuspended pellet, the supernatant of the first centrifugation and a suspension of 1 µM pure protein in 50 µL of storage buffer. After heating at 95 °C for 5 min, 10 µL of each sample was loaded onto Mini-PROTEAN gels (Bio-Rad) next to 3 µL of ladder (New England Biolabs) and separated at 120 V in SDS running buffer. The gel was stained using Coomassie dye (GelCode) for 1 h at room temperature with rocking, then washed in ultra-pure water until bands were clear. Gels were imaged with a Bio-Rad ChemiDoc MP imaging system.

### Immunofluorescence microscopy

RPE-1 cells were cultured in ibiTreat (Ibidi) dishes at 20-40% confluence and incubated overnight in the same conditions used for propagation. Cells were then infected, as above, and grown to 1, 3 or 5 dpi before fixation with 1% paraformaldehyde for 10 min at room temperature and storage in PBS at 4 °C.

Cells were permeabilised by incubation in absolute ethanol and then in 0.5% Triton X-100 in PBS, both at 4 °C for 30 min. After three washes with PBS, permeabilised cells were incubated in primary antibodies for 1 h at 37 °C or overnight at 4 °C. Polyclonal anti-Pal antibody (chicken, 1:10, target peptide KDKPEVEVTKDNKEEAYR; Davids) was used alongside a monoclonal human anti-TSA56 antibody (undiluted) with 1 mg/mL bovine serum albumin. Another three washes in PBS followed, before incubation in secondary antibodies for 1 h in the dark at 37 °C. Cells were labelled with DAPI (1:1000), anti-chicken IgY secondary antibody (goat, 488 nm excitation, 1:1000; Abcam) and anti-human IgG secondary antibody (goat, 647 nm excitation, 1:1000; Invitrogen).

Images were taken as a Z-stack using an Eclipse Ti2 confocal microscope (Nikon) and 100x/1.4NA oil lens. Imaging was performed according to the parameters listed in Table 6. Images were processed by maximum intensity projection and 2D deconvolution and analysed using ImageJ ^17^ and Fiji ^18^.

### Immunoblot analysis of anti-Pal antibody

Lysate from IPTG-induced *E. coli* expressing ΔN60 Ot_Pal was separated by SDS-PAGE. Proteins were transferred to a nitrocellulose membrane (BioRad) via wet transfer for 1h at 140mA. Membranes were blocked for 30 min in 2% w/v milk powder dissolved in PBS with 0.1% Tween (ThermoFisher Scientific) (PBST). They were then probed with primary antibody, either 1:1000 Mouse 6x-His Tag Monoclonal Antibody (ThermoFisher Scientific) or 1:10 Polyclonal anti-Pal antibody, in 2% w/v milk powder/PBST at 4°C overnight with rocking. The next day, the membrane was washed three times in PBST for 10 min, then probed with secondary antibody as appropriate – either 1:1000 Goat Anti-Mouse IgG Polyclonal Antibody (IRDye® 680RD) (LicorBio) or 1:1000 Goat Anti-Chicken IgY H&L (Alexa Fluor 647) (Abcam) – in 2% w/v milk/PBST for 3h at room temperature with agitation, protected from light. The membrane was then washed thrice more in PBS for 10 min each and stored in PBS. Bands were visualised with a Bio-Rad ChemiDoc MP imaging system.

### *In silico* analyses

AlphaFold 3 was used to produce protein models and PAE plots. Signal sequences were determined using SignalP 6.0 to allow removal of these from models^15^. Models were produced for Ot strain Karp (UniProt A0A2U3RMG0) residues 19-204 and *Rickettsia prowazekii* Madrid E (UniProt Q9ZCH2) residues 18-155.

The following specific strains were used in sequence analyses: *Chlamydia trachomatis* serovar D; *Wolbachia pipientis* wMel; *Orientia tsutsugamushi* Karp; *Escherichia coli* K12; *Caulobacter crescentus* NA1000; *Rickettsia prowazekii* Madrid E.

### Proteome-wide signal peptide classification and lipobox inspection

The *O. tsutsugamushi* Boryong proteome (Uniprot reference; TaxID 357244; Proteome ID UP000001565, downloaded 2025-08-19) was analysed with SignalP 6.0 ([v 6])^15^ using the “slow-sequential” models and organism = Other. The ‘Other’ organism group was chosen because *O. tsutsugamushi* does not fit cleanly into the Gram-positive/Gram-negative categories used by SignalP. For each protein, SignalP output reports a class label (OTHER, Sec/SPI, Sec/SPII [LIPO], Tat/SPI, Tat/SPII [TATLIPO], or Sec/SPIII) and a predicted cleavage site (CS) reported as positions *i–j* with an associated probability, where the mature sequence starts at *j*.

Sequences classified as Sec/SPII (LIPO) or Tat/SPII (TATLIPO) were retained for inspection. Canonical bacterial lipoproteins were defined a priori by the presence of a cysteine at position +1 relative to the predicted cleavage site. For each retained sequence, the residue at position *j* was examined directly in the primary sequence. Any sequence lacking Cys was not considered a canonical lipoprotein.

### HMM-Based Presence/Absence Profiling Across Proteomes

We downloaded 8 reference proteomes from UniProt (Table 2), plus an *O. tsutsugamushi* Karp proteome constructed from a curated in-house database based on genomes analysed in Giengkam et al^19^.

We profiled lipoprotein maturation, trafficking pathway and the *β*-barrel assembly machinery: *lgt, lspA, lnt, pal; lolA–E; bamA–E*. For each locus, three well-annotated seed sequences were compiled into one FASTA per locus (Supplementary Table 2). Seeds were aligned with MAFFT^20^ (v7.505, --auto) and one HMM per locus was built with HMMER (v 3.3.2, hmmbuild)^21^.

Each locus HMM was searched against every reference proteome with HMMER (hmmsearch, v3.3.2)^21^. Domain tables (--domtblout) were parsed to derive a binary presence/absence call and to record the best-scoring hit. A locus was considered present in a given proteome if at least one domain hit satisfied i-E ≤ 1×10^−^ □ and target coverage ≥ 0.50.

We computed the target coverage as: 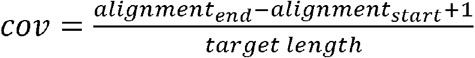 where *alignment_start* and *alignment_end* are the residue indices (1-based) of the first and last aligned positions in the target protein, and *target_length* is the full length of the target protein sequence.

Independently of these cutoffs, for each (species, locus) pair we also recorded the best domain hit, defined as the target sequence with the lowest independent E-value, and its associated coverage.

Some HMMER searches produced cross-hits to related protein families and false positives, which created inconsistencies in the initial presence/absence matrix (for example, unexpected absences, additional copies, or borderline hits near the score thresholds). The hmmsearch-derived calls were therefore used as a first-pass screen, and discrepant or borderline cells were re-evaluated using manual sequence analysis, structure-based methods and literature review.

## Acknowledgements

The authors are grateful to Shannon McKie at CIMR for assistance with setting up the liposome pelleting assays, and to members of the lab for support and discussions.

## Figure Legends

**Table 1. Summary of plasmids used**.

**Table 2. Reference proteomes used for HMM profiling**.

**Supplementary Figure 1. Signal peptide and lipobox predictions in AmiD, MlaA and RlpA**. Signal peptides were predicted using SignalP 6.0 (dark blue) and, where that was unsuccessful, DeepTMHMM-1.0 (light blue). Lipobox motifs are predicted using SignalP 6.0 and indicated by a red box (solid line), with the target cysteine residue shown in red. Incomplete lipobox motifs lacking a cysteine residue are shown with a dotted red line. The canonical lipobox motif is shown.

**Supplementary Figure 2. Immunoblot analysis of anti-Pal antibody**. Immunoblot demonstrating specific binding of Anti-6xHis (left) and Anti-Pal (right) antibodies to *E. coli* overexpressing Ot_Pal.

**Supplementary Table 1. Analysis of nine bacterial genomes to determine the presence or absence of lipoprotein-related genes**.

**Supplementary Table 2. Supplementary Table 2. UniProt seed sequences used to build locus-specific HMMs**.

