## Supplementary Figure 1 for "Complete loss of lipoprotein biosynthesis machinery in an obligate intracellular bacterium"

Lipobox motif [LVI]-[ASTVI]-[GAS]-[C]

| Group | Protein | Sequence |
| --- | --- | --- |
| AmiD (YbjR) | *Ec*_AmiD | MRRFFWLVAAALLLAGCAGEK |
| | *Cc*_AmiD | MSLSIIEAPSPNFDARKAVPDTVILHYTGMETGEAAIERLRD |
| | *Rc*_AmiD | MIVIDKTSYRAVGFDKRIKFLVLHYTQCDFKQSLD |
| | *Ot*_AmiD | MLYLPLLQLLQHVIVDNALVVNKSNVHQKSQLQTTISS |
| MlaA | *Ec*_MlaA | MKLRLSALALGTTLLVGCASSGTDQQG |
| | *Cc*_MlaA | MAPMRSRSIRLRDRHQSARPQGFRRHLAVGALAAAGLLAAASN |
| | *Rp*_MlaA | MKILIILSIILCSLFGRADLEYVDNDIYT |
| | *Ot*_MlaA | MRIIVAIIITCTTALMIKADTLSIKNSDNYKYTENIK |
| RlpA | *Ec*_RlpA | MRKQWLGICIAAGMLAACTSDDGQQQTVSVPQPAVCNGPIVEISG |
| | *Cc*_RlpA | MNQGIEQVSRFARRFCLMALAAFSLAACATPKYEVGRTVATSTGS |
| | *Rp*_RlpA | MIIMRNYKFKLEPLKQCQEAFKRYTVHSTALYIIALQANWSRRLIYKL |
| | *Ot*_RlpA | MNFFQKINMQLIGFAVSILLYNLPSICIAAQNNSICQINKQLLIM |
