## Supplementary figures and images for "Complete loss of lipoprotein biosynthesis machinery in an obligate intracellular bacterium"

### Supplementary Figure 2

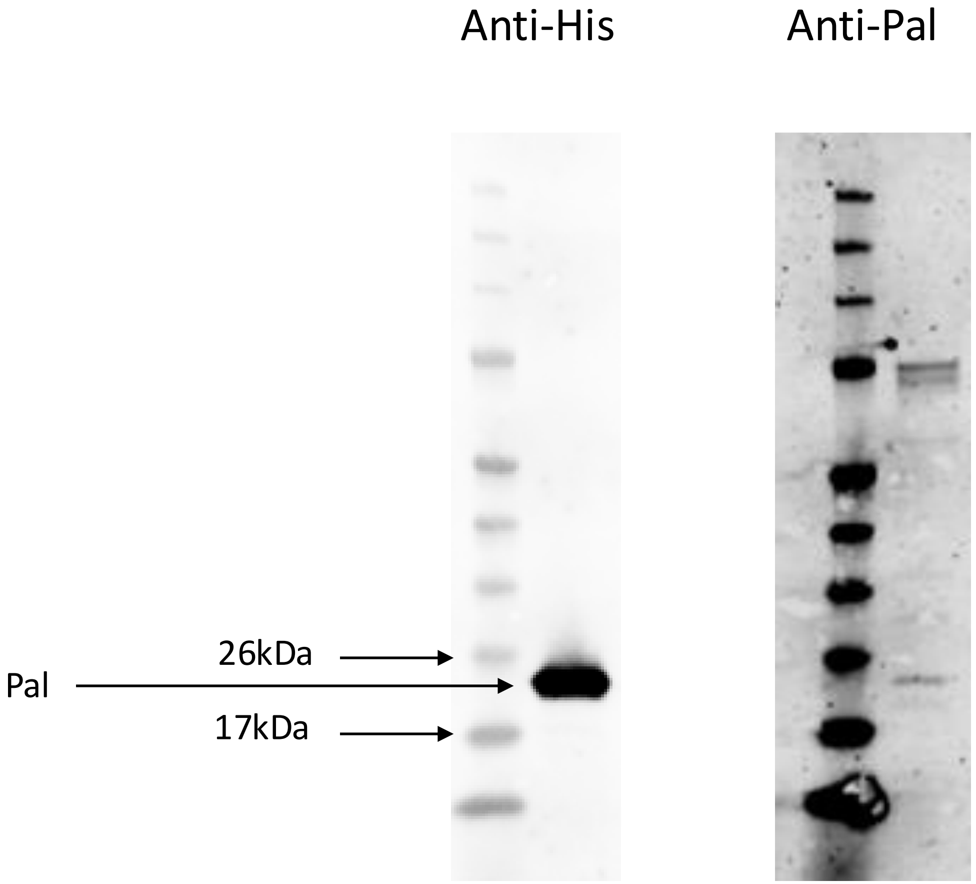
