## Supplementary material for "Complete loss of lipoprotein biosynthesis machinery in an obligate intracellular bacterium": Supp. Table 2

**Supplementary Table 2. UniProt seed sequences used to build locus-specific HMMs.**
Each locus-specific HMM was built from three UniProt seed sequences listed in this table.

| **Locus** | **UniProt Accession** | **Organism** |
| --- | --- | --- |
| bamA | P0A940 | *Escherichia coli* |
| bamA | D5CHY0 | *Enterobacter cloacae* |
| bamA | O25369 | *Helicobacter pylori* |
| bamB | P77774 | *Escherichia coli* |
| bamB | Q0A984 | *Alkalilimnicola ehrlichii* |
| bamB | F5ZAY7 | *Alteromonas naphthalenivorans* |
| bamC | P0A903 | *Escherichia coli* |
| bamC | C7RSI4 | *Accumulibacter regalis* |
| bamC | Q3SJU7 | *Thiobacillus denitrificans* |
| bamD | P0AC02 | *Escherichia coli* |
| bamD | Q50985 | *Neisseria gonorrhoeae* |
| bamD | O25930 | *Helicobacter pylori* |
| bamE | P0A937 | *Escherichia coli* |
| bamE | Q8EGW5 | *Shewanella oneidensis* |
| bamE | Q9K1F0 | *Neisseria meningitidis* |
| lgt | P60955 | *Escherichia coli* |
| lgt | Q9K266 | *Chlamydia pneumoniae* |
| lgt | P60966 | *Paenarthrobacter aurescens* |
| lnt | A0QZ13 | *Mycolicibacterium smegmatis* |
| lnt | Q8FJY4 | *Escherichia coli* |
| lnt | Q7VRA5 | *Blochmanniella floridana* |
| lolA | P61316 | *Escherichia coli* |
| lolA | Q9I0M4 | *Pseudomonas aeruginosa* |
| lolA | Q7VR38 | *Blochmanniella floridana* |
| lolB | P61320 | *Escherichia coli* |
| lolB | P42812 | *Pseudomonas aeruginosa* |
| lolB | Q476F7 | *Cupriavidus pinatubonensis* |
| lolC | P0ADC3 | *Escherichia coli* |
| lolC | P57062 | *Neisseria meningitidis* |
| lolC | Q8K9N8 | *Buchnera aphidicola* |
| lolD | P75957 | *Escherichia coli* |
| lolD | Q5LUD0 | *Ruegeria pomeroyi* |
| lolD | Q89KN0 | *Bradyrhizobium diazoefficiens* |
| lolE | P75958 | *Escherichia coli* |
| lolE | P44250 | *Haemophilus influenzae* |
| lolE | A0A5S9NPF1 | *Starkeya nomas* |
| lpp | P69776 | *Escherichia coli* |
| lpp | Q7CQN4 | *Salmonella typhimurium* |
| lpp | Q6D622 | *Pectobacterium atrosepticum* |
| lspA | P00804 | *Escherichia coli* |
| lspA | Q9HVM5 | *Pseudomonas aeruginosa* |
| lspA | Q661F0 | *Borrelia garinii* |
| pal | P0A912 | *Escherichia coli* |
| pal | P26493 | *Legionella pneumophila* |
| pal | P10324 | *Haemophilus influenzae* |
