## Supplementary material for "Complete loss of lipoprotein biosynthesis machinery in an obligate intracellular bacterium": Tables

| **Plasmid name** | **Construct name** | **Base plasmid** | **Composition of construct** |
| --- | --- | --- | --- |
| pGW1 | FL Pal | pET29b(+) | Pal sequence encoding residues 18-204, with C-terminal His-tag (GSG linker and His_6_), inserted under the T7 promoter. |
| pGW2 | ΔN60 Pal | pET29b(+) | Pal sequence encoding residues 61-204, with C-terminal His-tag (GSG linker and His_6_), inserted under the T7 promoter. |

**Table 1. Summary of plasmids used.**

| Strain | Accession | Proteome | Source |
| --- | --- | --- | --- |
| *Escherichia coli* | GCA_000005845.2 | UP000000625 | UniProt |
| Bacillus subtilis | GCA_000009045.1 | UP000001570 | UniProt |
| *Mycoplasma genitalium* | GCA_000027325.1 | UP000000807 | UniProt |
| *Borrelia burgdorferi* | GCA_000008685.2 | UP000001807 | UniProt |
| *Chlamydia trachomatis* | GCA_000008725.1 | UP000000431 | UniProt |
| *Caulobacter crescentus* | GCA_000022005.1 | UP000001364 | UniProt |
| *Wolbachia pipientis* | GCA_001752665.1 | UP000175679 | UniProt |
| *Rickettsia prowazekii* | GCA_000195735.1 | UP000002480 | UniProt |
| *O. tsutsugamushi* Karp | -- | -- | In-house ref^18^ |

*^18^O. tsutsugamushi* Karp in-house reference proteome curated based on Giengkam *et al.* (mSphere 8, e0026823, 2023)

**Table 2. Reference proteomes used for HMM profiling.**
